# Fish avoid visually noisy environments that reduce their perceptual abilities

**DOI:** 10.1101/2020.09.07.279711

**Authors:** Joanna R. Attwell, Christos C. Ioannou, Chris R. Reid, James E. Herbert-Read

## Abstract

The environment contains different forms of ecological noise that can reduce the ability of animals to detect information. Here we ask whether animals can adapt their behaviour to either exploit or avoid areas of their environment with increased dynamic visual noise. By immersing three-spined sticklebacks (*Gasterosteus aculeatus*) into environments with a simulated form of naturally occurring visual noise – light bands created by the refraction of light from surface waves termed caustic networks – we tested how such visual noise affected the movements, habitat use, and perceptual abilities of these fish. Fish avoided areas of higher visual noise, and achieved this by increasing their activity as a function of the locally perceived noise level, resulting in individuals moving away from noisier areas. By projecting virtual prey into the environment with different levels of visual noise, we found that the fish’s ability to visually detect prey decreased as visual noise increased. We found no evidence that fish increased their exploration (and decreased their refuge use) in environments with increased visual noise, which would have been predicted if they were exploiting increased visual noise to reduce their own likelihood of being detected. Our results indicate that animals can use simple behavioural strategies to mitigate the impacts of dynamic visual noise on their perceptual abilities, thereby improving their likelihood of gathering information in dynamically changing and noisy environments.

## Introduction

Animals live in inherently noisy environments, where noise can be defined as any environmental stimulus that interferes with the ability of animals to detect or respond to biologically meaningful cues (Brumm, 2013; Corcoran and Moss, 2017; Cuthill et al., 2017). Noise can take acoustic, chemical or visual forms, and can also vary dynamically in the environment. For example, wind-blown vegetation, turbidity (Chamberlain and Ioannou, 2019), and the refraction and scattering of light through water (Matchette et al. 2018; Matchette et al. 2020) can create backgrounds with dynamically changing illumination, while weather, traffic or other human activities can generate varying intensities of background acoustic noise (Lampe et al., 2012; Morris-Drake et al., 2016; Slabbekoorn and Ripmeester., 2008; Tasker et al., 2010; Vasconcelos et al., 2007). These forms of noise can reduce the likelihood of animals detecting information in their environment through two processes. First, by adding statistical error to the sensory modality being utilised, noise can make detection of a stimulus within that sensory channel more difficult due to masking effects. Alternatively, by distracting an animal, noise may limit its ability to detect or respond to information across sensory modalities (Morris-Drake et al., 2016). Through these processes, noise can reduce the ability of animals to communicate with conspecifics, (Fleishman, 1986; Herbert-Read et al., 2017; Lampe et al., 2012; Ord et al., 2007; Peters et al., 2007; Slabbekoorn and Ripmeester, 2008; Vasconcelos et al., 2007), detect moving targets (Matchette et al., 2018), forage efficiently (Azeem et al., 2015; Evans et al., 2018; Matchette et al., 2019; Party et al., 2013; Purser and Radford, 2011; Wale et al., 2013), and respond to predatory attacks (Morris-Drake et al., 2016; Wale et al., 2013), all of which have significant survival and fitness consequences for individuals.

Because noise can reduce the ability of animals to detect or respond to information in their environment, prey and predators may use behavioural strategies to either exploit or avoid noisy environments, thereby increasing their likelihood of detecting information, or avoid being detected themselves. For example, if attempting to remain undetected, some species may preferentially select noisier environments, or increase their exploration of the environment during times of increased environmental noise. Indeed, fathead minnows (*Pimephales promelas*), three-spined stickleback (*Gasterosteus aculeatus*), and larval pike (*Esox lucius*), show decreased anti-predator behaviour in more turbid water (Abrahams and Kattenfeld, 1997; Lehtiniemi et al., 2005; Sohel and Lindstrom, 2015), suggesting they may exploit times of high turbidity to avoid being detected by visual predators (although see (Chamberlain and Ioannou, 2019)). On the other hand, some species may attempt to avoid noisier environments as gathering information in those environments becomes more difficult. Indeed, some species of bats avoid areas of their environment with higher levels of acoustic noise (Bennett and Zurcher, 2013) and others spend more time foraging in areas with lower levels of acoustic noise (Schaub et al., 2008) (but see (Bonsen et al., 2015)). When avoidance of noisy areas is impossible, however, some species may adapt their behaviour to compensate for reduced information detection. Zebra finches (*Taeniopygia guttata*), for example, spend more time being vigilant in noisier acoustic environments, but this comes at the cost of decreased foraging rates (Evans et al., 2018). While animals’ behavioural changes to noise, and in particular acoustic noise, have been relatively well documented (Kunc et al., 2016; Shannon et al., 2016), whether animals adapt their behaviour in response to noise in other sensory channels, and in particular dynamic visual noise, has received far less attention. Moreover, whether these changes to behaviour represent adaptive behavioural decisions to exploit or avoid noisy environments remains unclear.

Shallow aquatic environments are particularly prone to one form of naturally occurring dynamic visual noise – *caustics*. Caustics are formed from the diffraction and refraction of light from the disturbance of surface waves that is projected through the water column onto the substrate below (Lock and Andrews, 1992). This form of dynamic illumination can reduce the likelihood of humans detecting a target on a computer animated display (Matchette et al., 2018), and can increase the latency of triggerfish (*Rhinecanthus aculeatus*) to attack a moving target (Matchette et al., 2020). Because caustics appear to reduce the ability of animals to detect or respond to information, animals could try to mitigate the impacts of, or exploit, these visually noisy environments. For example, species might avoid areas with higher levels of visual noise to increase the likelihood of detecting information themselves, or alternatively, could preferentially associate with visually noisy environments to reduce their own likelihood of being detected. Here we ask how visual noise affects the movements, refuge use and prey detection abilities of three-spined sticklebacks (*Gasterosteus aculeatus)*. Sticklebacks are a small (∼ 2 - 6 cm) fish found in both shallow marine and freshwater environments where caustics are prevalent. They feed on small insects, fish fry, and crustaceans by actively searching for prey among the substrate and in the water column, and are themselves predominantly predated by birds and larger fishes, such as pike (*Esox lucius*). To understand whether sticklebacks can exploit or avoid these visually noisy areas, we performed a series of three experiments. We first asked whether stickleback preferred to associate with visually noisy environments (potentially to reduce the likelihood of themselves being detected by their own predators), or avoid those areas (potentially to increase the likelihood of detecting their own prey). In this experiment, we also determined whether a potential preference was driven by the fish actively or passively choosing to avoid or associate with noisy environments by quantifying their movements in response to the locally perceived level of noise, that is, the level of visual noise that surrounded the fish. Second, we tested whether the fish increased or decreased refuge use in different levels of visual noise, predicting that if the sticklebacks were using visually noisy environments to avoid being detected by predators, they should use a refuge less frequently in higher levels of visual noise. Last, we tested whether visually noisy environments affected the ability of stickleback to detect prey in their environment by quantifying the likelihood that individual stickleback detected virtual prey in environments with different levels of visual noise.

## Methods

### Playbacks

For each experiment, we projected video playbacks of simulated caustics into an experimental arena to assess how different levels of visual noise affected the habitat choices, refuge use, and prey detection abilities of sticklebacks. These refracted patterns of light naturally vary in their spatial distribution, intensity, and velocity as a function of the water depth and the properties of the surface waves (Lock and Andrews, 1992). To produce the playbacks, we first generated a series of 600 images of caustic patterns using the software, Caustics Generator Pro (dualheights, 2018), cropped to an aspect ratio of 3840 by 2159 pixels (see software settings in Table S1). We created animations of these images in MATLAB 2018a where the images were stitched together sequentially (See Video S1). The animations were designed so that they could be looped without the caustics appearing to spatially or temporally ‘jump’. To create six different levels of visual noise, we wanted to ensure we manipulated only one property of the caustic patterns. For example, if we had manipulated the spatial fractal nature of the caustic patterns, this would have also changed the total light intensity within our animations. Therefore, we chose to manipulate the speed that the caustics moved, while maintaining their spatial properties. To do this, we created videos where the images looped through at different speeds, so that the slowest to fastest animation took 80, 40, 20, 10, 5 and 2.5 seconds, respectively, to complete a full loop. We classified faster moving playbacks as having higher levels of visual noise. We ensured that the speed at which the caustics moved in the arena, and the light levels in the arena, were representative of those found in nature (see Online Supplement & Figure S1). Furthermore, we chose to not include a treatment of a static projected image of the caustic patterns, as we were specifically interested in how different levels of visual noise affect behaviour, and note that static caustics are ecologically unrealistic and never occur in aquatic environments.

### Study subjects and experimental arena

Three-spined sticklebacks (n = 204) were caught from the river Cary in Somerton, Somerset, UK (51.069990 latitude, −2.758014 longitude), and we observed that caustics were present in the location that the fish were caught. Fish used in experiment one were caught in November 2017, and fish used in experiment two and three were caught in March 2019. All fish were in non-breeding condition when used in experiments. All fish were housed for at least two weeks before experimentation. The fish were housed in glass housing tanks (40 ⨯ 70 ⨯ 35 cm, height ⨯ length ⨯ width) on a flow-through re-circulating freshwater system with plastic plants and tubes for environmental enrichment. The fish were held at 14^*°*^C under a 11:13 hour light:dark cycle. Each tank housed approximately 40 individuals and fish were fed bloodworms once per day, six days per week.

The experimental tank for all three experiments consisted of a test arena (1.46 ⨯ 0.84 m, length ⨯ width) and a holding area (0.84 ⨯ 0.34 m, length ⨯ width) separated by white opaque plastic (Figure S2). Both sections were filled to a depth of 15 cm with water from the re-circulating freshwater system. Water was filtered within the tank using a Eheim classic 600 External Filter and chilled to between 14.2 and 15.2^*°*^C using a D-D DC-300 chiller. The holding area contained plastic plants and tubes for environmental enrichment. Water was changed in the experimental tank between each of the three experiments but not between individual trials. Prior to each day of trials, fish that would be used in the subsequent day were placed in the holding area overnight, allowing them to acclimate to the conditions of the tank. Fish were not fed for 24 hours prior to each experiment.

A camera (Panasonic HC-VX870) located 2.15 m above the centre of the test arena filmed the trials at 4K resolution (3840 ⨯ 2160 pixels) at 25 frames per second. The camera was controlled remotely using the Panasonic Image App operated by an experimenter outside the room. The experimenter was not present in the room while the trials were run. A BenQ W1700/HT2550 Digital Projector with 4K resolution operating at a 60 Hz vertical scan rate located 2.19 m above the arena (Figure S2) projected the playbacks into the arena. The projections were played using a Dell Inspiron 15 notebook connected to the projector via a 4K HDMI cable located outside the experimental room. The arena was surrounded by black-out curtains to minimise any external light source entering.

### Experiment 1 – Do fish prefer to associate with more or less visually noisy environments

To determine if fish tended to avoid or preferred to associate with more or less visually noisy environments, individual fish were presented with a binary choice, where on one side of the arena we projected one level of visual noise, and on the other side we projected a different level of visual noise. As there were six different levels of noise, this gave 15 possible combinations of noise pairings. We constructed six different playbacks, where each playback contained all 15 different noise pairings, played one after the other. Each choice (noise pairing) was presented for 320 seconds, with the total length of each playback equalling 80 minutes. Across the different playbacks (n = 6), each noise level was presented evenly on each side of the test arena to control for any potential side biases (Table S2). The ordering of the noise pairings within a playback were also assorted (see Table S2) between the six playbacks so there was no systematic bias in their ordering across the trials. For each trial, a single fish was exposed to one of these six playbacks. The order of the six playbacks was randomised within each day, with each playback being used a maximum of once per day.

For each trial (n = 48), an individual fish (5.4 cm ± 0.7 cm, mean ± SD) was moved from the holding area into the test arena and left to acclimate there for 10 minutes. During this period, the first frame of the playback was projected into the test arena as a static image, but the video playback was not started. After ten minutes, we started the video playback remotely. After each trial, the fish was removed and placed in a separate housing tank and fed. No fish were reused between trials.

Fish were tracked using an adapted version of DIDSON tracking software (Handegard and Williams., 2008) in MATLAB 2018a (Figure 1a). Because the fish were darker than the arena or moving projections, we took a grey-scale threshold of each frame to isolate the fish within the videos without requiring any background subtraction. Tracks were filtered to remove spurious tracking errors and smoothed using a rolling average of 12 frames (approximately half a second). In addition to plotting and manually inspecting the tracks for accuracy, we calculated that fish were tracked for 88.2% of all frames (see below). The tracking accuracy was not systematically affected by different levels of visual noise (see Online Supplement).

**Figure 1:**
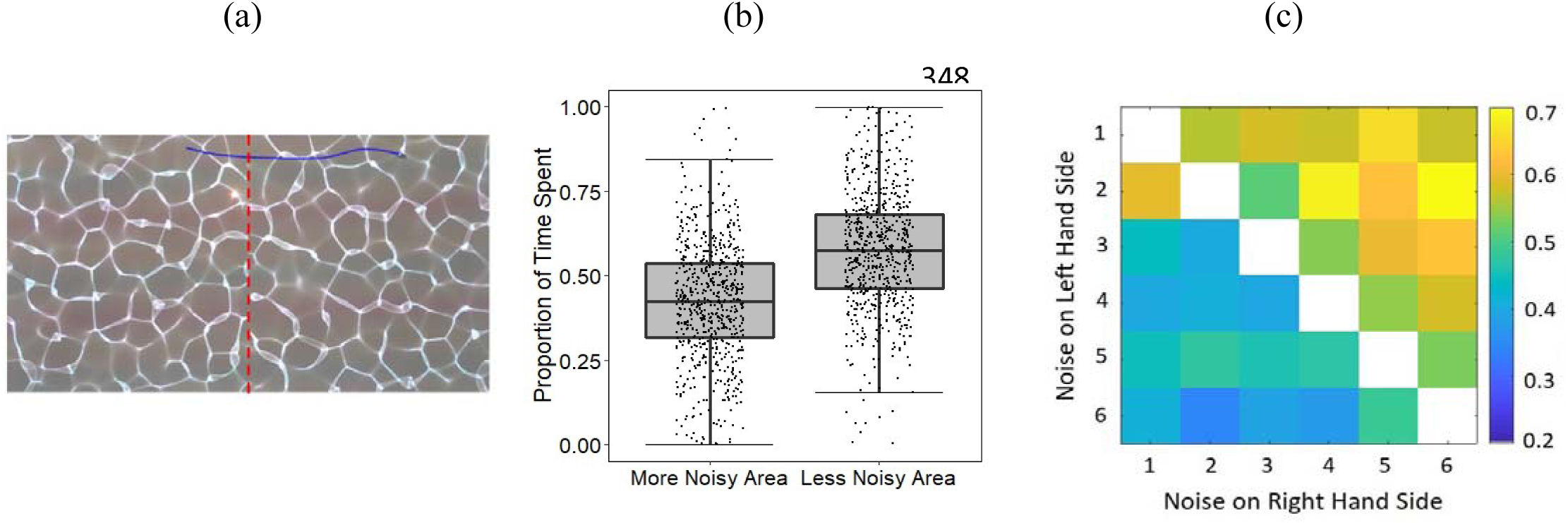
Experimental arena and choice test. a) Still image from a video of a choice trial depicting a section of the trajectory of a fish superimposed in blue. The red dotted line shows the virtual boundary between the two choice areas. b) Proportion of time the fish spent on the side of the arena with more or less visual noise (ignoring absolute differences in noise level). Fish spend more time on the side of the arena with less visual noise. The central line of each box shows the median value while the upper and lower lines of the box show the upper and lower quartiles of the data, with the whiskers extending to the most extreme data point within 1.5 × the interquartile range. Jittered dots represent raw data points. c) Matrix showing the mean proportion of time across trials (the heat) the fish spent on the left side of the arena where each cell represents the choice a fish was given between two levels of visual noise. Six represents the highest level of noise, and one the lowest. When the noise level was *lower* on the left side of the arena (upper-right corner of the plot), the fish spent *more* time on that side. When the noise level was *higher* on the left side of the arena (lower-left corner of the plot), the fish spent *less* time on that side. Note that 1 minus this matrix would give the proportion of time spent on the right-hand side of the arena.

From the trajectory data of each fish, we calculated the amount of time the fish spent on each side of the test arena in each paired choice. To do this, we used the *inpolygon* function in MATLAB 2018a to determine when a fish’s track was either on the left or right side of the arena. We then calculated the proportion of time that the fish spent in the noisier side of the arena for each noise pairing. We also calculated the amount of time the fish spent stationary, and the speed they adopted when they were moving, when in different levels of noise. To do this, the instantaneous speed of the fish was calculated as the displacement in their position between two consecutive frames. We defined a fish as being stationary when its speed was less than 2 mm s^−1^, informed by plotting a histogram of all the fish’s speeds across all trials (see Figure S3). The mean speed of a fish was calculated excluding the times when they were stationary (i.e. when speeds were *>* 2 mm s^−1^), and was calculated when the fish was on each side of the arena separately.

### Experiment 2 – Do fish use refuge more or less in increased levels of visual noise

To determine whether the fish were more or less likely to use a refuge in different levels of visual noise, two plastic plants (each 5 ⨯ 2 ⨯ 15 cm, length ⨯ width ⨯ height at the base) were placed as a refuge in the middle of the test arena (see Figure S4). For each trial (n = 48), an individual fish (4 ± 0.5 cm; mean standard length ± SD) was exposed to six different levels of visual noise, with each level of noise being projected throughout the entire arena (unlike Experiment 1), and each level of noise projected sequentially one after the other. To ensure that each noise level was presented in a different order within trials, we again created six playbacks where each playback contained each level of noise, but each noise level occurred in a different order within each playback (referred to as order-within-trial) in a Latin-square design (see Table S3). Each trial, therefore, consisted of six levels of noise, each presented for 320 seconds, with the total running length of the playbacks equalling 32 minutes. Every playback was presented once per day (between 9am and 5pm) and in a random order on different days. As in the choice tests, individual fish were moved from the holding area and placed in the test arena for 10 minutes acclimation time (while a static caustic image was projected into the arena) before the playback was started remotely. Experimented fish were removed and placed in a separate housing tank and later fed. Used fish were kept separately from unused fish and fish were not reused between trials. Fish that were used in the refuge experiment had not been used in the choice experiment.

We scored the amount of time (in seconds) the fish spent under the refuge during each level of noise. To do this, videos were imported into the software BORIS v. 7.9.15 (Friard et al., 2016), where we defined the fish to be under the refuge when any part of its body was under any of the fronds of the plastic plant (see Figure S4). Each fish could therefore be under the refuge for a minimum of 0 to a maximum of 320 seconds in each level of visual noise.

### Experiment 3 – Does visual noise affect the ability of fish to visually detect prey in their environment?

To determine if different levels of visual noise affected the ability of fish to detect prey in their environment, we created playbacks of the six different levels of visual noise (as in Experiment 2, played sequentially and throughout the entire arena), that included virtual prey that appeared and disappeared in random locations. In particular, the prey appeared as uniform red dots (similar to the virtual-prey experiments in (Ioannou et al., 2019; Duffield and Ioannou, 2017)) on top of the caustic patterns (see Video S2). When these playbacks were projected into the arena, each virtual prey appeared at a random location within the arena as a looming stimulus, increasing in size from 0 - 12.5 mm diameter within 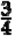of a second, maintaining 12.5 mm diameter for ∼ 1 second, and then shrinking to 6 mm before disappearing. Each prey was visible in the arena for a total of two seconds, moving on a correlated random walk at 7 cm *s*^−1^. Playbacks of the caustics and prey were again created in MATLAB 2018a.

Because the sticklebacks were fed on red bloodworm in their housing tanks, they were highly responsive to these red dots, often attempting to peck at them if in range. Therefore, the fish did not need to be trained to attack the virtual prey. However, the limited presentation time of each prey (2 s) was designed to allow the fish to detect and start swimming towards the prey, but reduce the likelihood of the fish sampling the prey, and hence learning that the prey was not edible. Due to the short presentation times, the arena being large, and the prey appearing in random locations, the prey would often appear far from the fish’s position. To increase the potential for the fish to detect the prey, therefore, each level of noise contained 50 individual prey presentations (300 presentations within a single trial), with four seconds between the end of one prey presentation and the start of another.

As in the refuge experiment, for each trial (n = 108), individual fish (3.3 ± 0.4 cm; mean standard length ± SD) were exposed to six levels of visual noise including the virtual prey. To control for order effects, six different playbacks were created where each playback contained each level of noise, with each noise level occurring at different times (order-within-trial) in the playbacks in a Latin Square design (Table S3). Controlling for order effects was particularly important here, as there was the potential for the fish to become habituated to the prey over time (although this did not occur – see below). In these playbacks, we also added transitions between the different levels of noise, so that the change in speed of the moving caustics between different noise levels was smooth. This involved creating animations that increased or decreased in speed from one noise level to another, which were subsequently placed between the respective animations of visual noise with the prey. Within these transition periods, no prey were projected. As in the other experiments, each level of noise lasted for 320 seconds, with each transition period lasting between 70 to 90 seconds. Each playback in total, therefore, lasted for ∼ 50 minutes and 45 seconds. Each playback was presented once per day and at different times of the day (between 9am and 5pm) on different days. As in the previous two experiments, individual fish were moved from the holding area to the test arena and allowed 10 minutes acclimation time. During this time, the lowest level of noise was projected into the test arena for ten minutes before transitioning into the start of the playback. Experimented fish were removed and placed in a separate housing tank and fed. No fish that were used in the choice or refuge experiments were used in this experiment and fish were not reused between trials.

Videos of the trials were manually inspected to determine whether the fish detected each of the virtual prey. A detection was defined as when there was a noticeable change in the speed or direction of the fish towards the prey (see Video S2 for examples). We quantified how many prey (out of a maximum of 50) the fish detected in each level of noise within each trial.

### Statistics

All statistics were performed in R 3.5.1 (R Core Team, Version 3.5.1). The package lme4 (Bates et al., 2015) was used for all mixed models. Assumptions for all linear mixed models (LMMs) were checked using standard diagnostic plots (QQ normal plots and residuals plotted against fitted values). Models were also checked for collinearity. Assumptions for all generalised linear mixed models (GLMMs) were checked using the DHARMa package (Hartig, 2019) including checking the dispersion and the distribution of the residuals. The full models were simplified by removal of non-significant terms. We used the anova function in R (R Core Team, Version 3.5.1) to compare pairs of models using the chi-squared statistic. The estimates and effect sizes (cohen’s D) are presented in Table S4. All R graphs were created using *ggplot2* (Wickham, 2016). All data used here are available in the dryad digital repository (doi:10.5061/dryad.rfj6q577x).

#### Experiment 1

To test whether fish spent more or less time in areas with more or less visual noise (regardless of absolute noise level), we subtracted 0.5 from the proportion of time they spent on the noisier side of the arena separately for each of the 15 noise pairings per trial. We tested whether the intercept of a linear mixed model (LMM) predicting those proportions, with trial included as a random effect, differed from zero (i.e. proportion of time on the noisier side – 0.5∼ 1 + (1 | Trial) in *lme4* nomenclature). We then asked whether the absolute difference in noise level within a choice affected the amount of time the fish spent on the noisier side of the arena. To do this, we calculated the difference between noise levels on each side of the arena for each noise pairing. For example, the difference between a choice of noise levels one and five was calculated as four. We then used an LMM to ask whether this difference could predict the proportion of time the fish spent on the noisier side of the arena. In this model, the difference in noise was treated as a discrete numeric variable, trial (fish identity) was included as a random intercept, and difference in noise level was included as a random slope.

In some choices (53 out of 720 choices) a fish did not visit both sides of the arena. These instances might not reflect, therefore, a true choice of the fish, as a fish could have been unaware of the noise level on the other side of the arena. Therefore, to test whether our results were robust to the removal of these instances, we performed the same analyses as described above, but only included data from when a fish had visited both sides of the arena during a choice. Our results did not qualitatively change when these cases were removed (see Online Supplement), and therefore we present results below including these cases.

The proportion of time that the fish spent in different levels of noise could result from fish adopting different movements as a function of the noise level they were in, and potentially the noise level on the other side of the arena. For example, if the fish adopted different speeds, or spent less time moving, this could result in the fish spending unequal amounts of time in each level of noise. To investigate this, we used LMMs to predict whether a fish’s speed, and in a separate model the time spent stationary, on the side of the arena the fish was in could be predicted based on the level of noise on each side of the arena (modelled as separate fixed effects: noise level on the side occupied by the fish and noise level on the unoccupied side). Mean speed and the proportion of time stationary were square root transformed due to a slight positive skew of these data. Order-within-trial (1-6) was also added as a fixed effect and trial was included as a random intercept along with noise level on the side occupied by the fish as a random slope.

Because a fish’s speed, and time spent stationary, were dependent on the noise level they were in (but not dependent on the noise level on the other side of the arena – see below), we asked whether the differences observed in speed and time spent stationary could solely explain the amount of time fish spent in the corresponding levels of visual noise. To do this, the mean speed that the fish adopted in different levels of noise, along with the proportion of time spent stationary, were added as covariates to the model investigating proportion of time spent on the side of the arena with more visual noise.

#### Experiment 2

We tested if the level of visual noise had a significant influence on the time the fish spent under refuge. We initially attempted to model the time spent in the refuge as a binomial response (time in versus time out of shelter), however, these models were over-dispersed. Therefore, due to the relatively bimodal distribution of the response variable (Figure S5a) we transformed the time spent in refuge into a binary response variable, where fish that spent over 50% of their time in the refuge were given a value of 1, and less than or equal to 50% a value of 0. This binary response variable was modelled using a GLMM with a binomial error structure. Noise level was included as a fixed effect (discrete numeric as before) along with order-within-trial, and trial (fish identity) added as a random effect. We did not include noise as a random slope in the refuge experiment because there was no effect of noise on the response variable.

#### Experiment 3

To test if the level of visual noise affected the likelihood that fish detected the prey, we used a GLMM with a Poisson family error structure. The response variable was the number of prey detections at each level of noise, and fixed effects were the level of visual noise and order-within-trial. Trial was added as a random intercept to account for the nonindependence of each noise level within a trial. Noise was not added as a random slope in this case due to the model not converging.

## Results

### Experiment 1 – Do fish prefer to associate with more or less visually noisy environments?

Fish spent more time on the side of the arena with less visual noise (Figure 1b) (LMM; *t*_47_ = −7.8, *p <* 0.001). Furthermore, the relative difference between the noise levels on each side of the arena affected the time the fish spent in the noisier side of the arena. As the relative difference in noise between the two sides of the arena increased, fish spent less time on the side of arena with more visual noise (Figure 1 c; LMM; _χ_^2^ = 9.77, *df* = 7, *p* = 0.002).

The fish’s movements were only affected by the level of visual noise on the side of the arena they were in. Fish moved faster (Figure 2a; LMM; _χ_^2^ = 52.3, *df* = 8, *p <* 0.001,), and spent less time stationary, when on the side of the arena with more visual noise (Figure 2b; LMM; χ2 = 90.8; *df* = 8, *p <* 0.001). The noise level on the other side of the arena (to that which the fish was on) did not affect the fish’s speed (Figure 2c; LMM; _χ_^2^ = 0.35, *df* = 8, *p* = 0.55), nor the proportion of time it spent stationary (Figure 2d; LMM; _χ_^2^ = 0.34, *df* = 8, *p* = 0.56). When these movement variables were added as covariates to the model, the difference in noise level between the two sides of the arena was no longer a significant predictor of the time the fish spent on the noisier side (LMM; _χ_^2^ = 1.61, *df* = 11, *p* = 0.20). The fish’s speed and its time spent stationary on the noisier side of the arena, therefore, could explain the proportion of the time spent on that side of the arena. In other words, how a fish adapted its movements to the locally perceived level of noise determined the amount of time it spent in that region.

**Figure 2:**
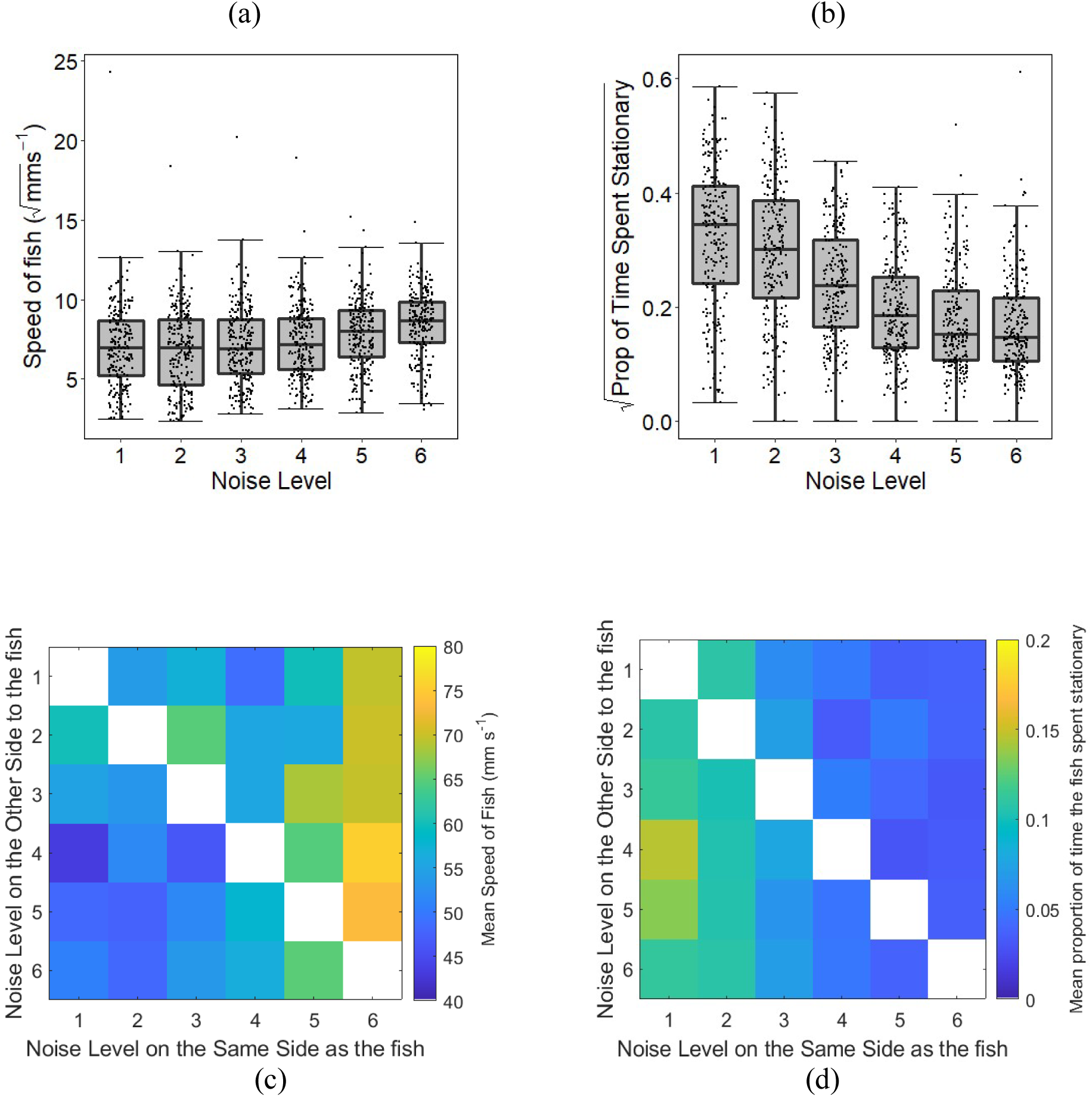
How the fish’s movements were affected by visual noise. a) Square root of fish’s mean speed as a function of the visual noise level they were in. Fish swam faster in more noisy areas. b) Square root of the proportion of time the fish spent stationary as a function of the noise level they were in. Fish spent less time stationary in more noisy areas. In a) and b) the central line of each box shows the median value while the upper and lower lines of the box show the upper and lower quartiles of the data. The whiskers extend to the most extreme data point within 1.5 × the interquartile range. Outliers are shown by the larger grey circles. Jittered points represent raw data points. c) Mean speed of fish (the heat) as a function of the noise level the fish was in (columns) and the noise level on the other side of the arena (rows). d) Mean proportion of time the fish spent stationary as a function of the noise level the fish was in (columns) and the noise level on the other side of the arena (rows). One – six corresponds to the lowest – highest levels of noise respectively. The presence of a trend from left to right, but not top to bottom, in these heat plots indicates that the fish moved faster, and spent less time stationary, in higher levels of visual noise, but the noise level on the other side of the arena did not affect their movements.

### Experiment 2 - Do fish use refuge more or less in increased levels of visual noise?

There was no evidence that the level of visual noise affected whether the fish spent the majority of time under the refuge or not (Figure S5b; GLMM; χ ^2^ = 1.75, *df* = 4, *p* = 0.19). However, fish did spend less time under the refuge as the trial progressed (GLMM; _χ_^2^ = 7.07, *df* = 4, *p* =0.0079).

### Experiment 3 - Does visual noise affect the ability of fish to visually detect prey in their environment?

Fish were less likely to detect the virtual prey in higher levels of visual noise (Figure 3; GLMM; _χ_^2^ = 156.6, *df* = 4, *p <* 0.01). There was no effect of the order in the trial on the number of detections by the fish (GLMM; _χ_^2^ = 0.003, df = 4, p=0.99); fish did not become less likely to detect the prey over the course of the trial.

**Figure 3:**
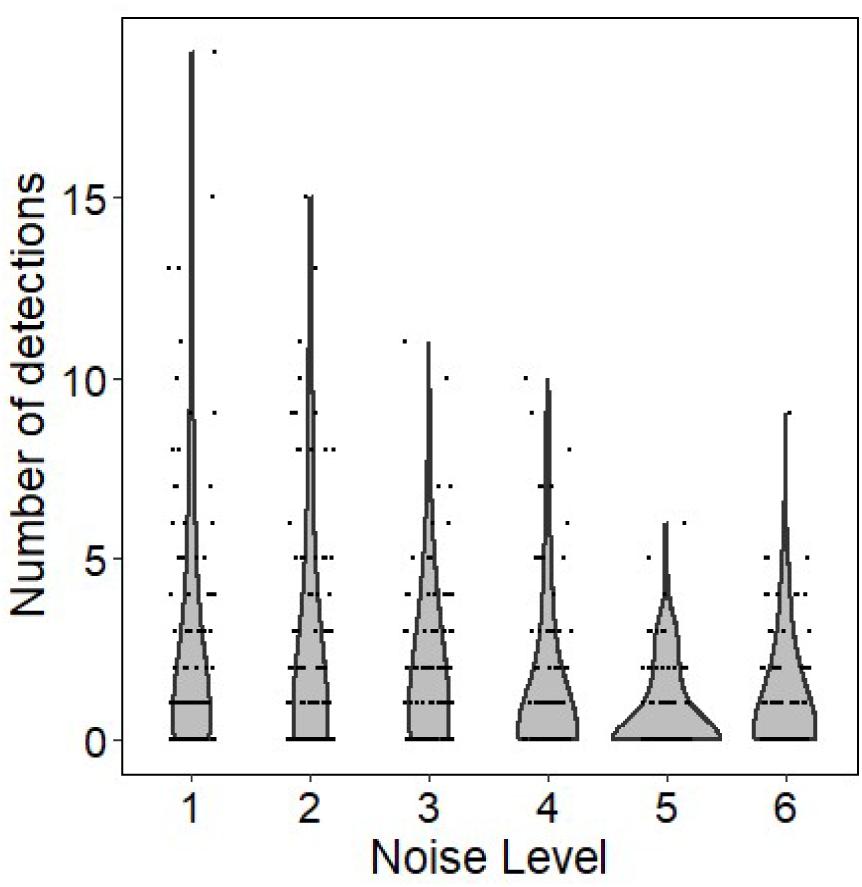
Virtual prey experiment. Number of times the fish detected the virtual prey out of the possible 50 prey presentations in each level of visual noise per fish (n = 108 fish in total). The violin represents a mirrored probability density function, and each black marker represents an individual data point jittered for clarity. Fish were less likely to detect the virtual prey in higher levels of visual noise.

## Discussion

Fish spent less time in areas with more visual noise, and this reduction in time could be attributed to how the fish adapted their movements in response to noise. In particular, fish increased their speed and spent less time stationary in areas with more visual noise. There was no evidence, however, that the level of visual noise on the other side of the arena affected their movements, suggesting that the fish were only responding to the level of noise in their local vicinity. While increases in speed and decreases in time spent stationary could be interpreted as the fish exploiting these environments to increase exploration during times of increased environmental noise, our second experiment provides evidence against this explanation. If fish were exploiting times of higher visual noise to avoid being detected themselves, we would have expected the fish to spend less time in the refuge in higher levels of visual noise (as refuge use is a key measure of risk taking in sticklebacks (Bevan et al., 2018)). In fact, we found no evidence that fish altered their risk-taking behaviour depending on noise level. Further, it is unlikely that fish were increasing their activity in more noisy areas to match their swim speed with the movements of the caustics, as the optical flow produced by the caustics did not move in a consistent direction (see Figure S1a & Supplementary Video 1). Instead, we suggest that fish use a simple mechanism, namely increasing their speed and activity, to avoid areas of their environment with higher levels of visual noise. This mechanism will lead fish to passively move out of these areas, and towards regions of the environment with lower visual noise. Indeed, similar mechanisms have been proposed for how groups of animals collectively track resources in their environment (Berdahl et al., 2013; Hein et al., 2015), providing a simple, yet effective mechanism to move towards or away from particular regions of the environment.

Our final experiment provided further support that these visually noisy environments should be avoided by sticklebacks, with increased visual noise reducing their ability to detect prey. Fish were less likely to detect the virtual prey in these environments, consistent with other systems where humans, chicks and triggerfish took longer to detect prey on backgrounds with dynamic visual noise as opposed to static controls (Matchette et al., 2018, Matchette et al., 2019, Matchette et al. 2020). Because animals have finite time and energy reserves, and limited attention (Cuthill et al., 2019), they are expected to make optimal foraging decisions that increase the rate or efficiency at which they gather resources (Schoener, 1971; Stephens and Krebs, 1986; Ydenberg et al., 1994). This allows them to devote more time and energy to other fitness-related activities (Pianka, 1988; Schmid-Hempel, 1991). Much like how animals choose foraging patches based on their profitability (Krebs, 1979; Milinski, 1979; Milinski, 1987), we might expect animals to selectively choose where to forage in their environment based not only on the resource profitability of a patch, but also considering the likelihood of detecting those resources given the perceptual constraints imposed by that environment. Indeed, there is large natural variation in both the temporal and spatial distribution of caustics in the aquatic environment, as well other forms of environmental noise. Caustics, for example, are prominent on clear, sunny days in intermediate water depths (0.25 - 5 m), but are reduced in deep waters and on overcast days with minimal surface waves. Such variation may lead foragers to select habitats based on the environmental noise determined by the local ecological conditions (e.g. Bennett and Zurcher (2013); Schaub et al. (2008)).

While fish avoided visually noisier environments, we may also expect individuals’ behavioural responses to environmental visual noise to vary as a function of other factors. For example, sticklebacks have been shown to both decrease (Sohel and Lindstrom, 2015) or increase (Ajemian et al., 2015; Chamberlain and Ioannou, 2019) their anti-predator behaviour and refuge use in more turbid water (a form of static visual noise). This suggests their response to visual noise could vary depending on context or state. Indeed, fifteen-spined stickleback (*Spinachia spinachia*), are less risk averse when hungry, but when partially satiated, choose less productive areas where they can spend more time being vigilant (Croy and Hughes, 1991). In our experiments, we did not feed the fish for 24 hours prior to the experiments to induce exploratory behaviour, and to promote search and targeting of the virtual prey. However, stickleback are both predators and prey, hence it would be valuable to test if the fish also avoid visually noisy environments when they are satiated, or when the level of risk in the environment is greater. Indeed, we might expect animals to choose noisier areas of the environment when satiated or when faced with greater risk. While not measured here, animals might also switch to relying on other sensory modalities in visually noisy environments when their vision is compromised (Partan, 2017; Suriyampola et al., 2018). For example, female three-spined sticklebacks rely more on visual cues when choosing a mate in clear water, but in turbid water, where vision is compromised, they rely more on olfactory cues (Heuschele et al., 2009). When habitats consistently differ in their ecological noise, this could have sweeping effects on populations. For example, grey squirrels from acoustically noisy urban habitats (*Sciurus carolinensis*) respond more to visual alarm signals than squirrels from rural habitats, which instead rely more on acoustic alarm signals (Partan et al., 2010). Hence animals from different populations may show different behavioural responses, highlighting the need for cross-population comparisons. We might also expect animals to rely more heavily on social information in sensory demanding environments. Indeed, animals in groups often benefit from pooling imperfect estimates of the world around them in order to make more accurate decisions collectively (Berdahl et al., 2018; Dall et al., 2005; Ioannou, 2017; Ward et al., 2011). Using playbacks of caustic visual noise, it would be possible to test how reliance on social information changes when individuals are exposed to an ecologically relevant form of environmental noise that reduces their visual perceptual abilities.

## Conclusion

Our results demonstrate that natural forms of ecological noise reduce the likelihood of animals detecting information in their environment. In response, animals can adapt their behaviour to avoid noisy areas, ultimately increasing their likelihood of gathering information, and thereby compensating for the negative impacts environmental noise has on their perceptual abilities.

## Acknowledgments

J.R.A was supported by a NERC GW4+ CDT PhD studentship NE/L002434/1. J.E.H.-R. was supported by the Whitten Lectureship in Marine Biology and a Swedish Research Council grant number 201804076. C.C.I was supported by Leverhulme Trust Grant RPG-2017-041 V. C.R.R was supported by an Australian Research Council Discovery Early Career Researcher Award grant number DE190-10-15-13. The authors would like to thank Andy Radford, Innes Cuthill, Sam Matchette and Nick Scott-Samuel for useful discussions and advice. Thanks are also given to the staff at the Animal Services Unit at Bristol University and in particular Peter Gardiner for all his help and advice with caring for the fish.

