## Supplementary document for "Fish avoid visually noisy environments that reduce their perceptual abilities"

**Supplementary Material for:** **Fish avoid visually noisy environments that reduce their perceptual abilities**

### Ensuring playbacks of caustics were consistent with natural conditions

We first ensured that the speed of the moving light bands fell within the natural range of wave speeds that occur in nature. To measure this, we recorded the playbacks as they were projected into the arena using a camcorder (Panasonic HC-VX870) at 25 frames per second and 4K resolution located 2.15 m above the centre of the arena. We also recorded a video of a static frame of the playbacks. These seven videos (one for each playback and the static frame) were then imported into MATLAB (2018a), where we used an optical flow analysis to measure the spatial and temporal dynamics of the projected caustics using the function *opticalFlowLK*. This function detects the speed and direction of displaced pixels between frames in a video by giving displacement vectors of seemingly moving regions (Figure S1a). We then converted these displacement vectors into real world displacements, using the known pixel to mm conversion ratio, determined with a calibration board. We calculated the speed of displacement (in mm s^−1^) across all regions within the arena in the six different playbacks separately.

Across all playbacks, the distribution of speed of the displaced pixels across the arena showed a strong peak close to 2 - 3 mm s^−1^ with heavy right tails (Figure S1b). The median speed of the top ten percent of the displacement vectors increased as a function of the speed of the playbacks, with the median displacement of the slowest playback being 13.5 mm s^−1^ and the fastest playback being 109.9 mm s^−1^ (Figure S1c). These speeds are consistent with the natural surface wave speeds that occur in shallow aquatic habitats (Francis, 1951).

To ensure the light intensity was consistent across the different playbacks, we projected the same frames (n = 36 frames) from playbacks of the lowest noise level, an intermediate noise level, and highest noise level into the arena and measured the average lux readings at five different locations in the arena for 10 seconds. This gave a total of 180 lux readings for each of the noise levels. A HOBO MX2202 was used to take the lux readings and was controlled from the HOBO mobile app. The reader was setup to store a reading every second. We tested whether these lux readings differed between the noise levels using a generalised linear model fitted with a negative binomial error structure using the glm.nb function from the MASS package (Venables and Ripley, 2002) in R (R Core Team, Version 3.5.1). This was due to the strong positive skew and over dispersion in the data. Assumptions of the model were checked using the standard diagnostic plots in R by plotting the residuals vs fitted values and by checking the dispersion using the blmeco package in R (Korner-Nievergelt et al., 2015). There was no significant difference between the light intensities of the lowest (94.4 lux; IQR = 213.0), medium (95.3; IQR=192.3) or highest noise levels (95.3; IQR = 182.6) (Figure S1d; GLM; z = -0.17, p = 0.86), confirming that our playbacks had equivalent average light intensities across noise levels.

### Tracking accuracy in different levels of noise

To ensure our tracking accuracy was not systematically affected by the level of visual noise in the choice experiment we calculated the lengths of continuous tracked segments (i.e. unbroken periods when the fish were tracked) in each level of visual noise. We then tested whether the tracked segment length changed as a function of noise level using a linear mixed model (LMM). The segment length was log transformed due to a strong positive skew. The segments for each side of the tank per choice per trial were given a unique ID (for each trial there were 15 sets of choices between 2 levels of noise meaning each trial had up to 30 unique IDs). This meant that all segments within each choice could be grouped together and accounted for a random factor in the model. Noise level was used as the explanatory variable and the unique ID within trial was added as a random intercept. The median tracked segment length was found to be 3 seconds (IQR = 7.32) long and the mean was 7.48 seconds (SD = 14.76). Although these tracked segment lengths are low, the average tracking accuracy across all trials and noise levels was 88.2%, hence tracked segments were only broken for relatively short periods of time. Noise level did not affect the length of the tracked segments, and hence would not have affected the interpretation of our results (LMM; *Estimate* = 1.01, *_χ_*^2^ = 1.78, *df* = 5, *p* = 0.18).

### Experiment 1 – Supplementary results

Here we present the results for the choice experiment only including those choices where the fish visited both sides of the arena. The results did not qualitatively change when these cases were removed. Fish spent more time on the side of the arena with less visual noise (LMM; *t*_45_ = −8.4, *p <* 0.001). Furthermore, the relative difference between the noise levels on each side of the arena affected the time the fish spent in the noisier side of the arena. As the relative difference in noise between the two sides of the arena increased, fish spent less time on the side of arena with more visual noise (LMM; *_χ_*^2^ = 7.79, *df* = 7, *p* = 0.005).

**Supplementary tables and figures**

Table S1: Parameter settings used in Caustics Generator pro to create the caustics

| Parameter settings | values |
| --- | --- |
| Depth | 5 |
| Intensity | 0.05 |
| Amplitude filter | 1.36 |
| Frequency filter | 1.5 |
| Time filter | 40.06 |

Table S2: Order that the different levels of noise were presented within each playback for Experiment 1 (choice). The letter above each pair of columns refers to the name of the playback. The numbers refer to the lowest (1) to highest (6) level of visual noise.

Order within	a	b	c	d	e	f

trial	Left	Right	Left	Right	Left	Right	Left	Right	Left	Right	Left	Right

| 1 | 5 | 1 | 3 | 2 | 2 | 1 | 2 | 4 | 4 | 3 | 5 | 2 |
| --- | --- | --- | --- | --- | --- | --- | --- | --- | --- | --- | --- | --- |
| 2 | 4 | 3 | 1 | 4 | 6 | 4 | 6 | 1 | 2 | 6 | 3 | 6 |
| 3 | 6 | 2 | 6 | 5 | 3 | 5 | 5 | 3 | 5 | 1 | 4 | 1 |
| 4 | 1 | 2 | 1 | 5 | 5 | 6 | 2 | 5 | 5 | 6 | 6 | 5 |
| 5 | 3 | 5 | 6 | 2 | 1 | 4 | 6 | 3 | 4 | 2 | 2 | 4 |
| 6 | 4 | 6 | 3 | 4 | 2 | 3 | 1 | 4 | 3 | 1 | 3 | 1 |
| 7 | 3 | 6 | 1 | 6 | 1 | 3 | 6 | 4 | 3 | 5 | 3 | 2 |
| 8 | 5 | 2 | 2 | 4 | 6 | 2 | 3 | 2 | 1 | 2 | 5 | 4 |
| 9 | 4 | 1 | 5 | 3 | 4 | 5 | 1 | 5 | 4 | 6 | 6 | 1 |
| 10 | 4 | 2 | 5 | 2 | 4 | 3 | 6 | 5 | 4 | 1 | 3 | 4 |
| 11 | 3 | 1 | 6 | 4 | 2 | 5 | 3 | 4 | 2 | 5 | 2 | 6 |
| 12 | 5 | 6 | 1 | 3 | 1 | 6 | 2 | 1 | 6 | 3 | 1 | 5 |
| 13 | 4 | 5 | 3 | 6 | 4 | 2 | 5 | 4 | 5 | 4 | 4 | 6 |
| 14 | 2 | 3 | 4 | 5 | 6 | 3 | 2 | 6 | 6 | 1 | 5 | 3 |
| 15 | 1 | 6 | 2 | 1 | 5 | 1 | 1 | 3 | 2 | 3 | 1 | 2 |

Table S3: Order that the different levels of noise were presented within each playback for the refuge and virtual prey experiments. The letter above each of the columns refers to the name of the playback (n = 6 different playbacks).

| Order within trial | a | b | c | d | e | f |
| --- | --- | --- | --- | --- | --- | --- |
| 1 | 2 | 4 | 1 | 6 | 3 | 5 |
| 2 | 3 | 1 | 5 | 4 | 6 | 2 |
| 3 | 1 | 5 | 4 | 3 | 2 | 6 |
| 4 | 6 | 2 | 3 | 1 | 5 | 4 |
| 5 | 4 | 6 | 2 | 5 | 1 | 3 |
| 6 | 5 | 3 | 6 | 2 | 4 | 1 |

| Explanatory Variable | Main Response variable | Estimate | Cohen D |
| --- | --- | --- | --- |
| Choice Experiment |  |  |  |
| Proportion time noisy side -0.5 | *Intercept model* | -0.08 | NA |
| Proportion time noisy side | Noise difference | -0.02 | 2.14 |
| Fish swim speed | Noise on side of fish | 0.09 | 1.88 |
| Fish proportion of time stationary | Noise on side of fish | -0.001 | -2.70 |
| Fish swim speed | Noise other side to fish | -0.0002 | 1.86 |
| Fish proportion of time stationary | Noise other side to fish | -0.001 | -2.73 |
| Proportion time noisy side | Noise difference, Swim speed, Proportion of time stationary | 0.0066 | 3.31 |
| Refuge Experiment |  |  |  |
| Time in refuge | Noise level | 0.22 | 1.77 |
| Time in refuge | Order-within-trial | -0.45 | 1.77 |
| Virtual Prey Experiment |  |  |  |
| Number of prey detections | Noise level | -0.23 | -0.85 |
| Number of prey detections | Order-within-trial | 0.0003 | -0.85 |

Table S4: Estimates and Cohen’s D for each of the statistical tests

(a)

(b)

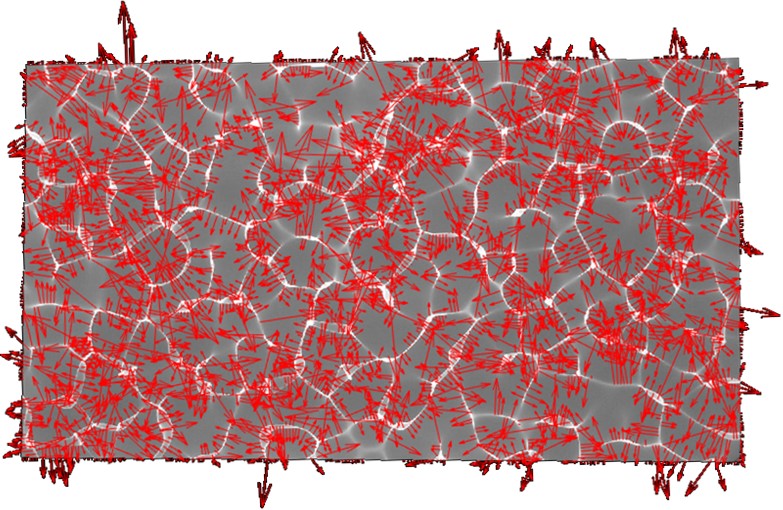

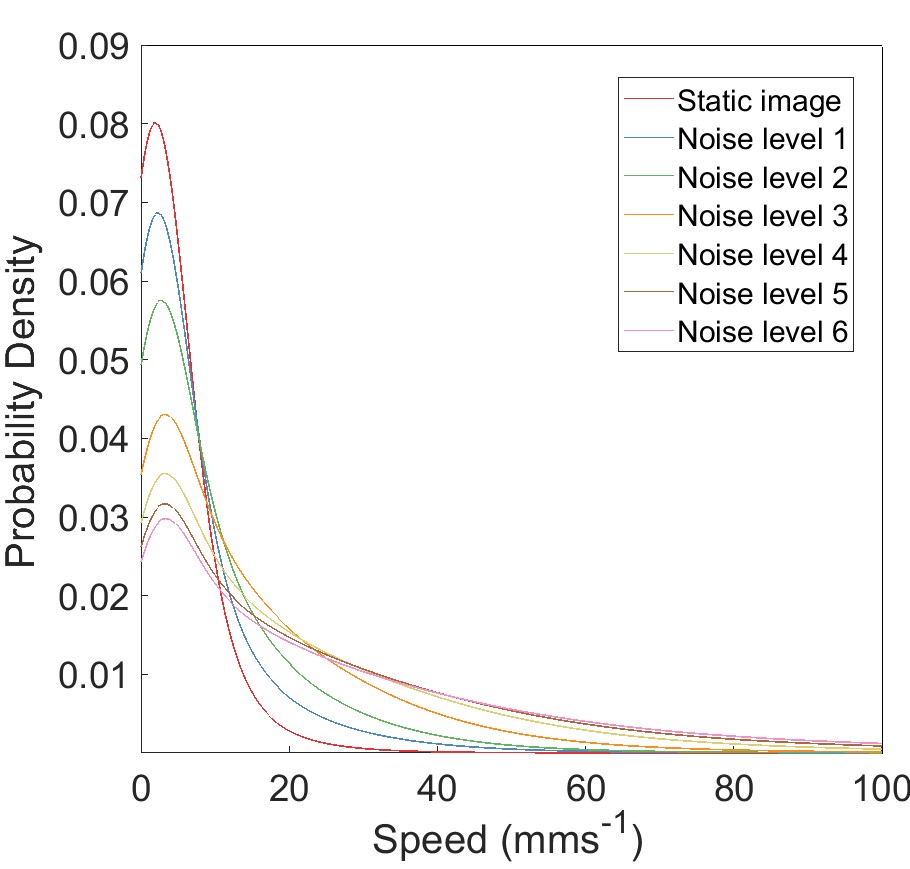

(c)	(d)

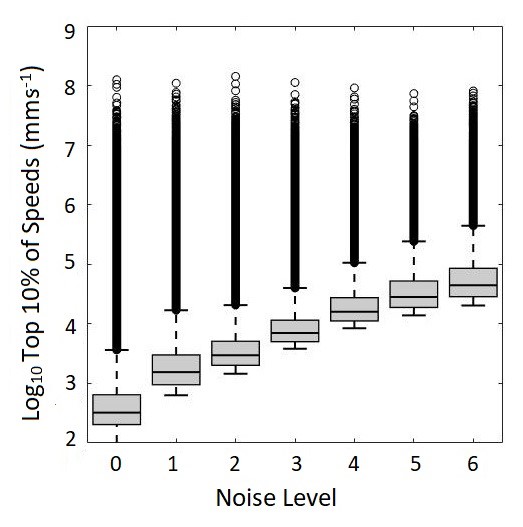

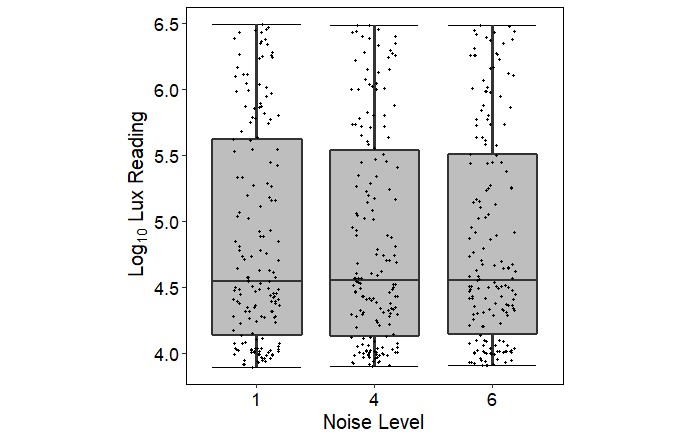

Figure S1: Quantifying the displacement of pixels and light intensities across the different playbacks. (a) Displacement vectors of regions within an image that were determined to have moved between two subsequent frames. For plotting purpose, vectors were decimated by a factor of [10 10] and scaled by a factor of 10 (b) Probability distribution of the speed of displacement vectors as a function of a static image and six different playbacks. (c) Boxplot showing the distribution of the top 10% of the speeds of displacement vectors as a function of the static image and six different playbacks. (d) Log lux readings taken across multiple frames in different locations for the lowest, medium and highest levels of noise. In c) and d) the central line of each box shows the median value while the upper and lower lines of the box show the upper and lower quartiles of the data, with whiskers extending to the upper and lower values not considered outliers (grey circles), points outside 1.5 × the interquartile range. Jittered points represent raw data points.

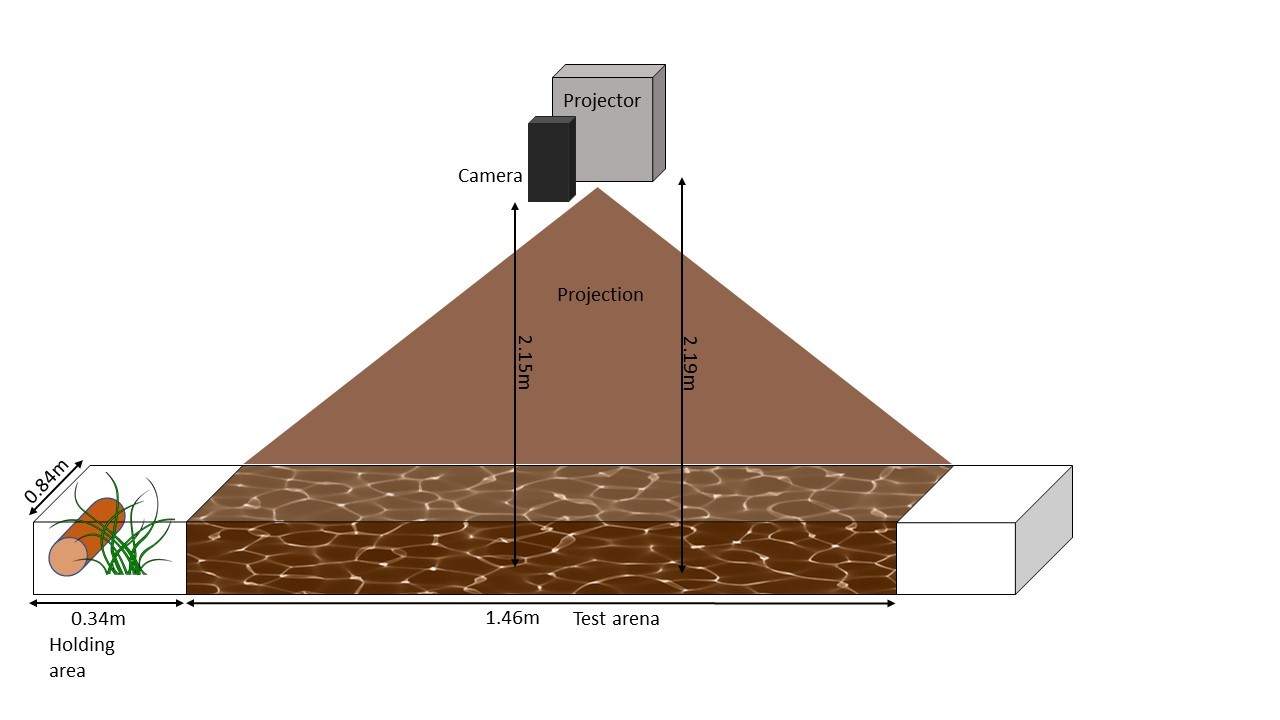

Figure S2: Schematic of the experimental set-up. The fish were kept in the holding area adjacent to the test arena overnight before being tested the next day.

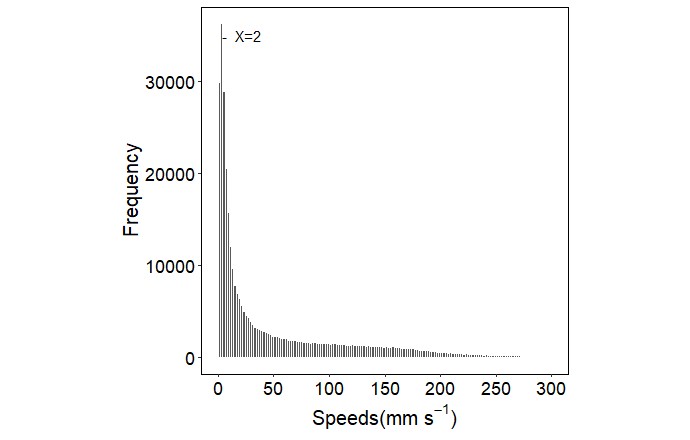

Figure S3: Distribution of all fish’s instantaneous swim speeds across all choice experiment trials. The line marked at X = 2 is to indicate the cut off, below which the fish was classified as being stationary.

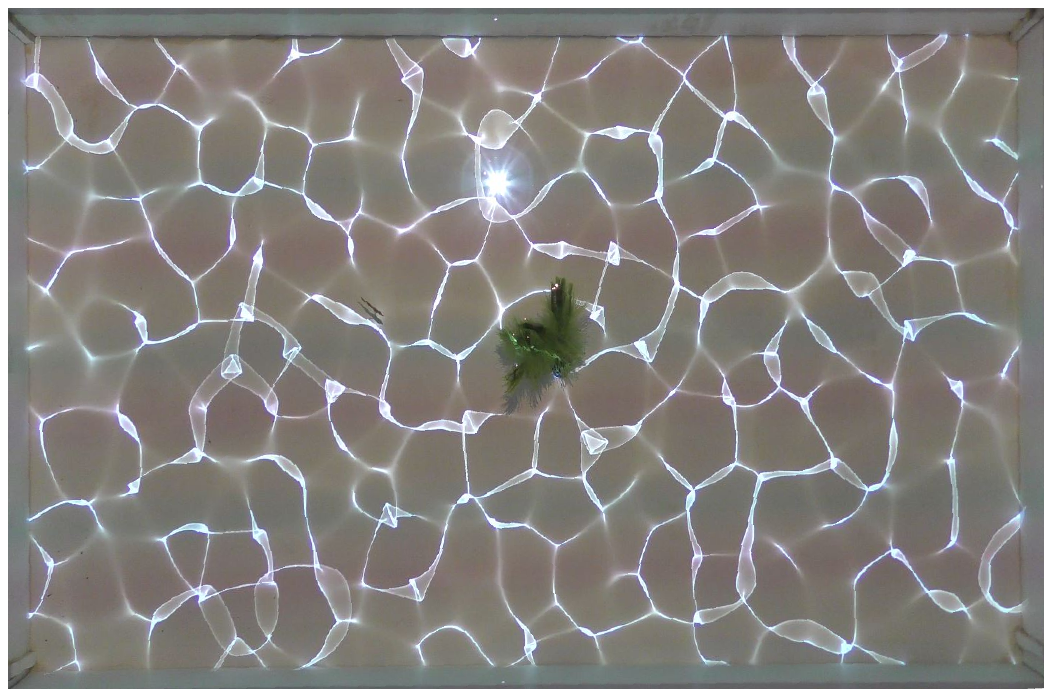

Figure S4: Setup of the refuge experiment with two plastic plants located next to each other in the middle of the tank. The size of each of the plants measured 5 x 2 x 15 cm (length x width x height), but as seen from the image, the fronds of the plastic splayed out when submerged.

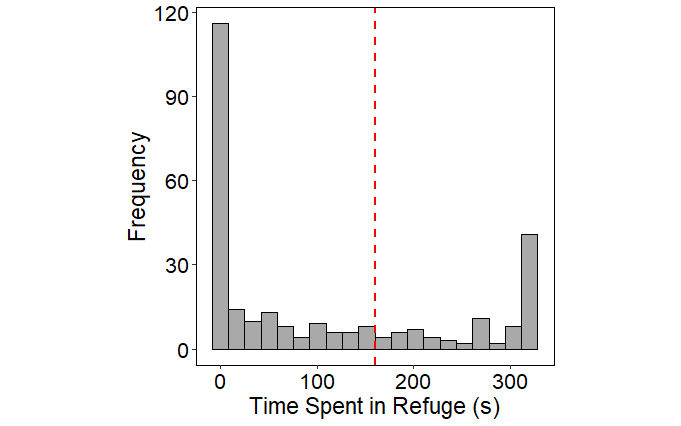
 (a)	(b)

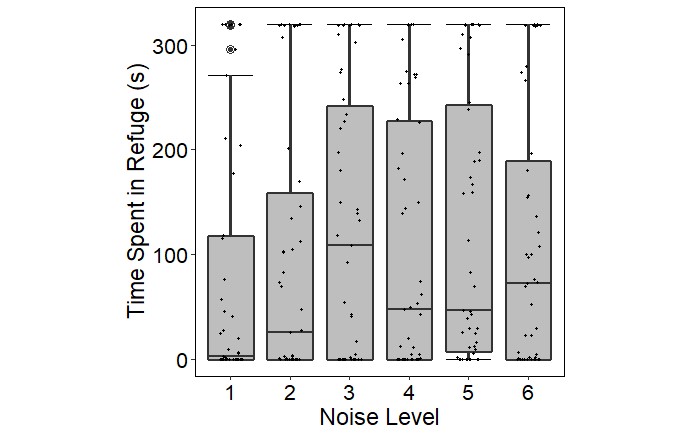

Figure S5: (a) Distribution of the time the fish spent in the refuge (max 320 seconds) for all levels of noise across all experiment trials. The red dashed line at x = 160 indicates the cut off, below which we classified fish as spending the majority of time out of the refuge (0), or above which we classified the fish as spending the majority of time in the refuge (1). (b) Amount of time the fish spent in the refuge as a function of noise level (however, note this was analysed using a binomial GLMM as defined in (a)). The time spent in the refuge was not affected by the noise level. The central line of each box shows the median value while the upper and lower lines of the box show the upper and lower quartiles of the data, with whiskers extending to the upper and lower values 1.5 × the interquartile range. Jittered points represent raw data points.

Springer, New York.
